# Depletion of the AD risk gene *SORL1* selectively impairs neuronal endosomal traffic independent of amyloidogenic APP processing

**DOI:** 10.1101/2020.01.06.896670

**Authors:** Allison Knupp, Swati Mishra, Refugio Martinez, Jacquelyn E. Braggin, Marcell Szabo, Dale W. Hailey, Scott A. Small, Suman Jayadev, Jessica E. Young

## Abstract

The *SORL1* gene encodes for the protein SorLA, a sorting receptor involved in retromer-related endosomal traffic. Many *SORL1* genetic variants increase Alzheimer’s disease (AD) risk, and rare loss-of-function truncation mutations have been found to be causal of late-onset AD. *SORL1* is expressed in neurons and glia of the central nervous system and loss of *SORL1* has been reported in AD tissue. To model the causal loss-of-function mutations, we used CRISPR/Cas9 technology to deplete *SORL1* in human induced pluripotent stem cells (hiPSCs) to test the hypothesis that loss of *SORL1* contributes to AD pathogenesis by leading to endosome dysfunction. We report that loss of *SORL1* in hiPSC-derived neurons leads to early endosome enlargement, a cellular phenotype that is indicative of ‘traffic jams’ and is now considered a hallmark cytopathology AD. We validate defects in neuronal endosomal traffic by showing decreased localization of amyloid-precursor protein (APP) in the trans-Golgi network (TGN), and increased localization of APP in early endosomes, a site of APP cleavage by the β secretase BACE1. Microglia, immune cells of the CNS, which play a role in AD pathology also express *SORL1*. We therefore tested and found no effect of *SORL1* depletion on endosome size or morphology in hiPSC-derived microglia, suggesting a selective effect on neuronal endosomal trafficking. Finally, because BACE1 dependent APP fragments can cause endosome enlargement, we treated *SORL1* deficient hiPSC-derived neurons with BACE1 inhibitors and demonstrate that endosome enlargement occurs independent of amyloidogenic APP fragments. Collectively, these findings clarify where and how *SORL1* links to AD. Moreover, our data, together with recent findings, underscores how sporadic AD pathways that regulate endosomal trafficking, and autosomal-dominant AD pathways that regulate APP cleavage, independently converge on the defining cytopathology of AD.

## Introduction

Alzheimer’s disease (AD) is the most common neurodegenerative disorder in the elderly affecting nearly 40 million people worldwide[1]. Currently, there is no treatment that alters disease progression. Recent therapeutic designs have focused on the main neuropathologic hallmarks of AD, accumulations of amyloid beta (Aβ) in senile plaques and abnormally phosphorylated tau protein in neurofibrillary tangles. The nearly universal failures of these trials to date, the vast majority of which have focused on removing or modulating Aβ, argues that other cellular pathways should be mechanistically studied for therapeutic development. Both genetics and pathology point to endosomal abnormalities and dysfunction as an early pathway in AD pathogenesis[2-5]. In particular, the *SORL1* gene, which encodes the sorting protein SorLA, is highly relevant, being associated with both late-onset and early-onset forms of AD[4, 6-9]. SorLA, first identified as a neuronal sorting receptor[10, 11], is expressed in nearly all cell types of the central nervous system[12] and has multiple roles in endocytic sorting[13-17], retromer-dependent retrograde trafficking[18], and regulating amyloid precursor protein (APP) processing[4, 10, 19, 20]. SorLA expression decreases in late-onset AD[21-23], and protein coding variants identified in early-onset AD families may lead to functional defects in the sorting of Aβ in cells[24]. Rare loss-of-function truncation mutations have been found to be causal of late-onset AD[7, 25]. We previously evaluated *SORL1* activity in human induced pluripotent stem cell (hiPSC)-derived neurons from AD patients and controls[20]. In these cells, *SORL1* expression induction with neurotrophic factors and its subsequent effect on neuronal Aβ peptides can be impacted by the presence of AD-associated risk variants.

Due to its role as a sorting receptor and because it may be decreased in AD, we hypothesized that *SORL1* deficiency would impact endosome pathology and, by default, trafficking of cargo in the endo-lysosomal network. To evaluate this hypothesis, we generated *SORL1* deficient hiPSC lines using CRISPR/Cas9 genomic editing. We examined endosomal size in two cell types differentiated from these hiPSCs--neurons and microglia. We also tested whether inhibiting amyloidogenic processing of APP by inhibiting β-secretase (BACE1) modulated endosome enlargement in hiPSC-derived neurons. In this study, we report that loss of *SORL1* by itself induces enlarged endosomes in hiPSC-derived neurons and this phenotype is not altered by BACE1 inhibition. We also observe that *SORL1* deficiency alters APP localization within the neuronal endosomal network. Interestingly, *SORL1* loss does not induce endosome enlargement in hiPSC-derived microglial-like cells, suggesting cell-type specific differences in this early AD cytopathology. Taken together, our data suggest that loss of the known AD risk gene *SORL1* induces early AD cytopathology in neurons, and that while it impacts trafficking of APP, the endosomal pathology occurs in an amyloid-independent manner. We observe important differences in two different cell types in endosome pathology underscoring the complexity of this cellular pathway in the central nervous system.

## Results

### *SORL1* depletion in human iPSC-derived neurons leads to enlarged early endosomes

We hypothesized that depletion of *SORL1* in human neurons would allow us to investigate early features of AD that may involve endosomal network dysfunction. We established isogenic *SORL1* KO and WT hiPSC lines using CRIPSR/Cas9 technology. We targeted exon 6 of the *SORL1* gene, inducing indels that disrupt the reading frame, leading to complete loss of SorLA protein (*SORL1* KO). (**Supplemental Figure 1**). In neurons differentiated from the *SORL1* KO hiPSC lines, we quantified staining of endogenous EEA1 and Rab5, (markers of early endosome morphology) using blinded, unbiased confocal microscopy. We observed significantly increased fluorescence intensity from EEA1 stained puncta (**Figure 1A-B**) and Rab5 stained puncta (**Supplemental Figure 3)** in *SORL1* KO neurons. Western blot analysis showed no change in total EEA1 protein between WT and *SORL1* KO neurons **(Supplemental Figure 2)**, suggesting that the increased fluorescence intensity is due to enlarged or fused early endosomes. We quantified endosome size and binned populations based on ≥ 0.5μm or <0.5 μm diameter distributions (**Figure 1C**). We observed significantly more endosomes greater than or equal to 0.5 μm in in *SORL1* KO neurons (**Figure 1C**), and a statistically significant increase in the mean EEA1 puncta area over the population of early endosomes as a whole (**Figure 1D**). Interestingly, we also noticed significantly enlarged endosomes in neural progenitor cells from the *SORL1* KO lines (**Supplemental Figure 3**) lines suggesting that loss of *SORL1* impacts endosome morphology early in the neural lineage, after hiPSCs are driven to neuroectoderm. Finally, we also tested an shRNA against *SORL1* in WT neurons and again observed significantly enlarged endosomes (**Supplemental Figure 3**), suggesting that an acute reduction of *SORL1* also leads to this phenotype.

**Figure 1.**
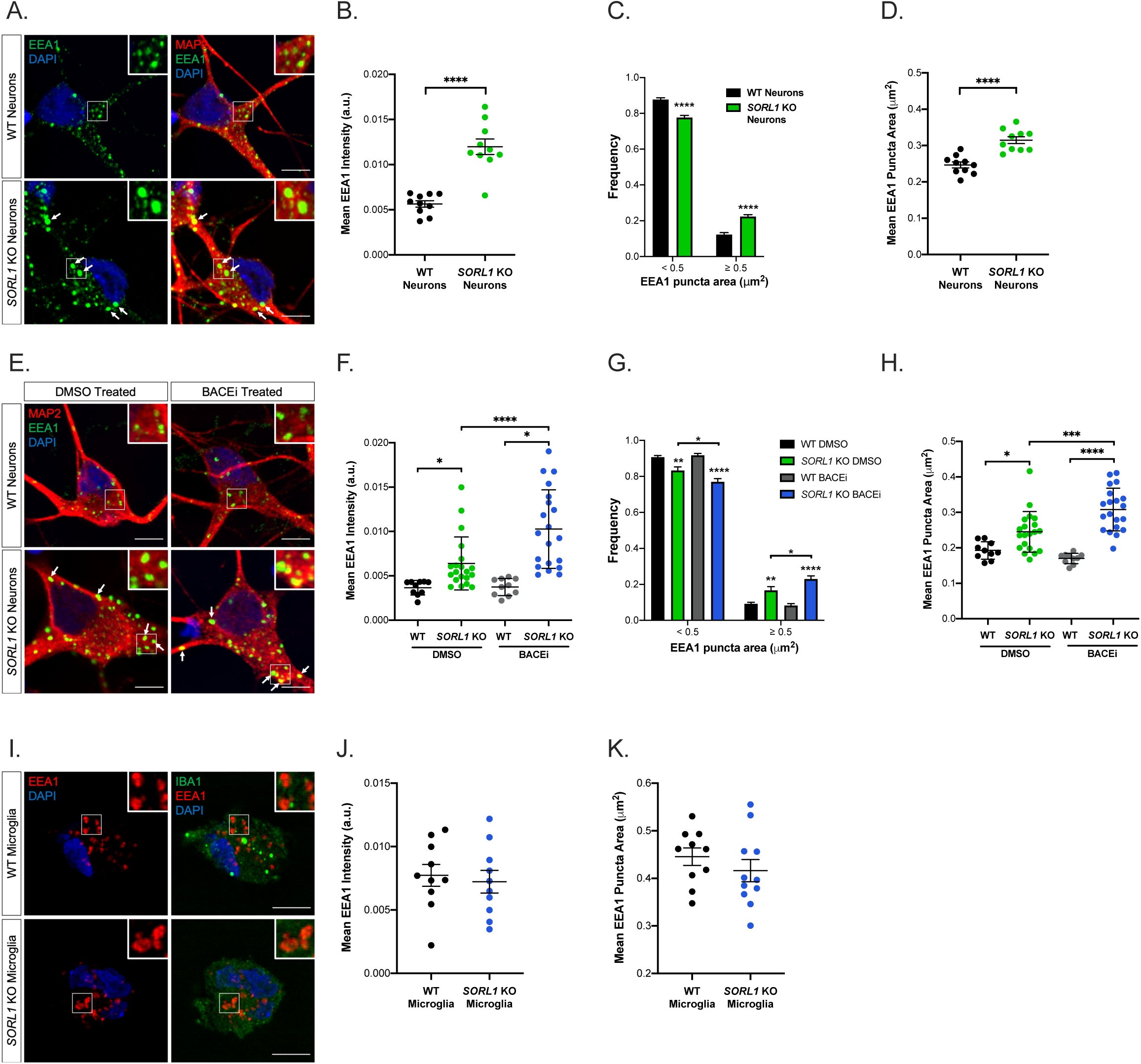
Depletion of *SORL1* leads to enlarged early endosomes in hiPSC-derived neurons but not microglia, independent of amyloidogenic APP processing. A. Representative immunofluorescence images of wild-type and *SORL1* KO neurons stained with EEA1 (green), MAP2 (red), and DAPI (blue). White boxes indicate regions of interest (ROIs) magnified in insets. Arrows indicate enlarged EEA1+ puncta. Scale bars are 5μm. B. Quantification of immunofluorescence images demonstrates increased EEA1 puncta fluorescence intensity in *SORL1* KO neurons compared to wild-type. n=10 images (18-22 cells). C. Size distribution of EEA1+ puncta in *SORL1* KO neurons compared to wild-type. n=6 images (18-22 cells) D. Quantification of mean EEA1+ puncta area in WT compared with *SORL1* KO neurons. N=10 images (18-22 cells). E. Representative immunofluorescence images of wild-type and SORL1 KO neurons treated with either DMSO or BACE1 inhibitor for 72 hours. Neurons were stained with EEA1 (green), MAP2 (red), and DAPI (blue). White boxes indicate regions of interest (ROIs) magnified in insets. Arrow indicate enlarged EEA + puncta. Scale bars are 5μm. F. Quantification of immunofluorescence images demonstrates increased mean EEA1+ puncta intensity in BACE1 inhibitor treated *SORL1* KO neurons compared to DMSO treated controls. n=10-20 images (42-58 cells). G. Size distribution of EEA1 puncta in *SORL1* KO neurons compared to wild-type with BACE1 inhibitor treatment. n=10-20 images (42-58 cells). H. Quantification of immunofluorescence images demonstrates enlarged EEA1 puncta area in BACE inhibitor treated *SORL1* KO neurons compared to DMSO treated controls. n=10-20 images (42-58 cells). I. Representative immunofluorescence images of wild-type and *SORL1* KO microglia stained with EEA1 (red), Iba1 (green), and DAPI (blue). White boxes indicate regions of interest (ROIs) magnified in insets. Scale bars are 5μm. J. Quantification of immunofluorescence images demonstrates no significant change in mean EEA1+ puncta intensity in *SORL1* KO microglia compared to wild-type. n=10 images (10-14 cells). K. Quantification of immunofluorescence images demonstrates no significant change in EEA1 puncta area in *SORL1* KO microglia compared to wild-type. n=10 images (10-14 cells). All values represent mean +/- SEM. Normally distributed data (B, C, D, H, J, K,) were analyzed by parametric statistical tests. Non-normally distributed data (F, G) were analyzed by non-parametric statistical tests. *p<.05, **p<.01, ***p<.001, ****p<.0001 by two-tailed unpaired T-test (B, C, J, K); by two-way ANOVA (D, H); or by Kruskal-Wallis test (F, G)

### Enlarged early endosomes in the context of *SORL1* depletion occur independent of amyloidogenic APP processing

Recent work has shown that neurons with introduced FAD mutations also show enlarged endosome morphology that is FAD gene-dose dependent[26]. In Kwart et al.’s study, endosome enlargement was dependent on β-secretase processing of APP. To test whether amyloidogenic processing of APP was also contributing to early endosome enlargement in the context of *SORL1* loss, we treated *SORL1* KO and isogenic WT neurons with an inhibitor of BACE1 (BACE1i), effectively reducing both Aβ peptide levels in our neurons (**Figure 2H**). Interestingly, in BACE1i treated neurons, we observed an even stronger increase in mean EEA1 intensity (**Figure 1 E-F**) and no amelioration of early endosome enlargement (**Figure 1 G-H)**. There was no difference in endosome size in WT cells treated with BACE1i as compared to DMSO treatment (**Figure 1 F).** This data suggests the mechanism by which loss of *SORL1* expression induces endosome enlargement is different from how FAD mutations impact the endosomal network, and is independent of amyloidogenic APP processing.

**Figure 2.**
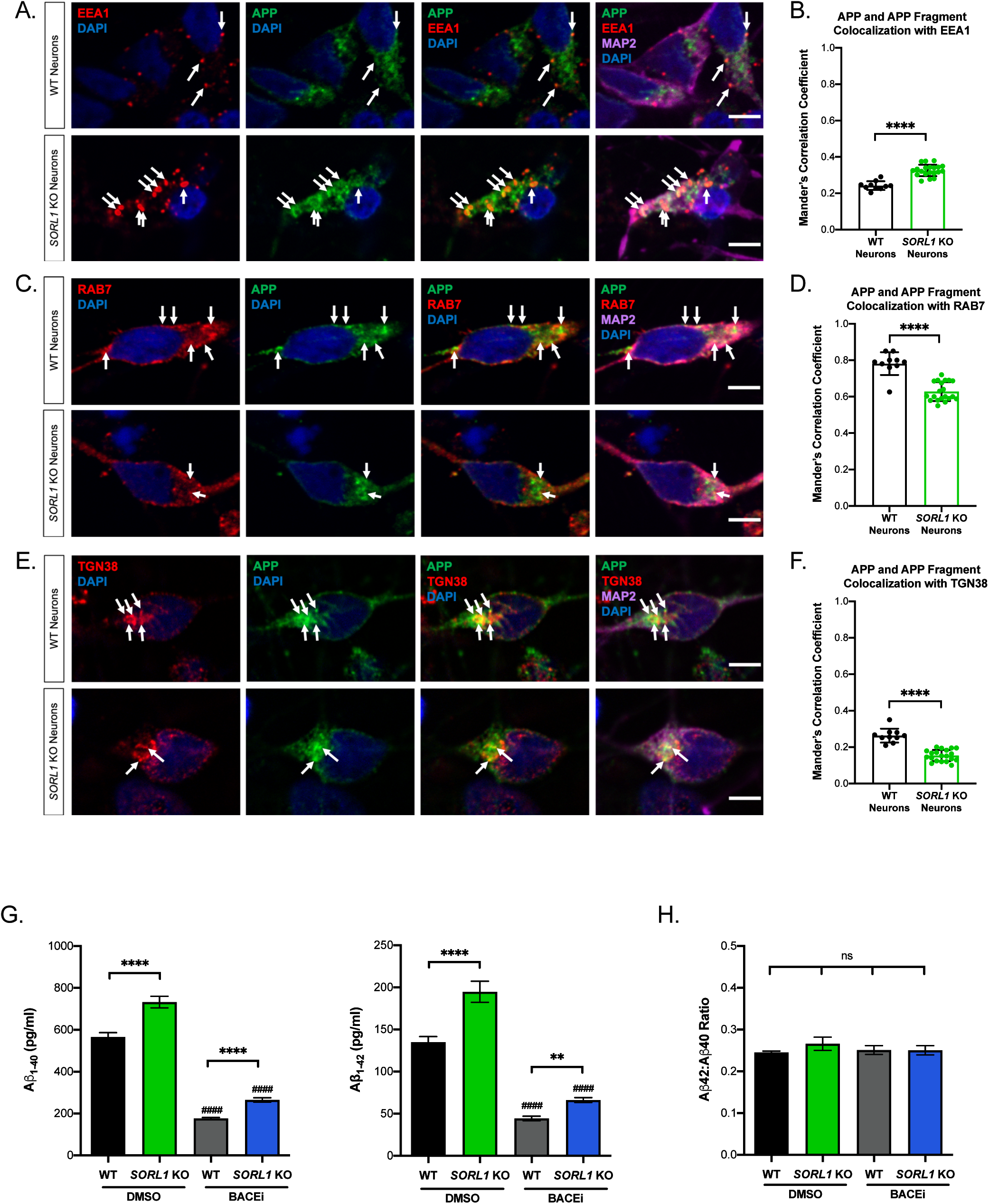
*SORL1* depletion in hiPSC-derived neuron alters APP localization in the endosomal network and increases amyloidogenic processing of APP. A. *SORL1* depletion results in APP accumulation in EEA1+ early endosomes. Representative immunofluorescence images of wild-type and *SORL1* KO neurons stained with EEA1 (red), APP C-terminal antibody (green), MAP2 (red), and DAPI (blue). Arrows indicate colocalization of EEA1+ puncta with APP and APP fragments. Scale bars are 5μm. B. Quantification of colocalization of EEA1 with APP, n=10-20 images. C. *SORL1* KO neurons show reduced colocalization of APP and APP fragments with Rab7+ puncta. Representative immunofluorescence images of wild-type and *SORL1* KO neurons stained with Rab7 (red), APP C-terminal antibody (green), MAP2 (red), and DAPI (blue). Arrows indicate colocalization of Rab7 puncta with APP and APP fragments. Scale bars are 5μm. D. Quantification of colocalization of Rab7 with APP, n=10-20 images. E. *SORL1* KO neurons show reduced colocalization of APP and APP fragments with TGN38+ puncta. Representative immunofluorescence images of wild-type and *SORL1* KO neurons stained with Rab7 (red), APP C-terminal antibody (green), MAP2 (far-red), and DAPI (blue). Arrows indicate colocalization of Rab7 puncta with APP and APP fragments. Scale bars are 5μm. F. Quantification of colocalization of TGN38 with APP, n=10-20 images. G. *SORL1* KO neurons have higher levels of Aβ_1-40_ and Aβ_1-42_ peptides than WT cells, indicated by asterisks (*). BACE1 inhibition significantly reduces these peptides in both genotypes as compared to their DMSO controls, indicated by hashmarks (#). In the presence of BACE inhibition *SORL1* KO neurons still have increased levels of Aβ_1-40_ and Aβ_1-42_ peptides, indicated by asterisks (*). H. Neither *SORL1* KO nor BACE1 inhibition changes the ratio of Aβ_1-40_ and Aβ_1-42_ peptides. All values represent mean +/- SD. All normally distributed data were analyzed by parametric statistical tests. *p<.05, **p<.01, ***p<.001, ****p<.0001 by two-tailed, unpaired T-test (B, D, F); or by two-way ANOVA (G, H).

### Enlarged early endosomes in *SORL1* deficient cells are present in hiPSC-derived neurons, but not microglial-like cells

AD-associated risk genes and their biological pathways may function differently between unique central nervous system cell types. Indeed, recent work using hiPSC-derived, gene-edited cells demonstrates that the strongest AD genetic risk factor, APOE ε4, impacts different cellular AD phenotypes in a cell type-specific manner[27, 28]. Microglia, the innate immune cells of the CNS, also express *SORL1*[12] but the functional role of *SORL1* in microglia is undefined and functionality of the endocytic network is likely very different in these highly phagocytic cells compared with neurons, which are professional secretory cells. We differentiated *SORL1* KO iPSCs to microglial-like cells using a previously published protocol[29] (**Supplemental Figure 4**) and analyzed endogenous EEA1 staining using similar protocols and analysis as for the hiPSC-derived neurons. Surprisingly, we did not observe any differences in early endosome fluorescence intensity (**Figure 1 I-J**) or size (**Figure 1 K)** when we compared WT and *SORL1* KO microglial-like cells. Microglia are derived from mesoderm/hematopoietic lineages while NPCs and neurons are derived from neural ectoderm. Our findings suggest that the endosomal trafficking and sorting functions of *SORL1* may depend on cell lineage and that *SORL1*-dependent early endosome pathology specific to neuronal cells.

### Loss of *SORL1* in hiPSC-derived neurons alters APP trafficking and processing in the endosomal network

Enlarged endosomes are indicative of endocytic traffic jams and may delay the proper maturation and progression of vesicles and cargo for processing and degradation[30]. One well-characterized cargo of *SORL1-*dependent trafficking is APP[18]. We quantified the localization of APP in various compartments of the endo-lysosomal network in *SORL1* deficient neurons using confocal microscopy. We observed an increase in colocalization of APP with EEA (**Figure 2 A-B**) and a decrease in colocalization of APP with TGN38, a marker for the trans-Golgi network (**Figure 2 C-D**), demonstrating reduced retrograde transport of APP in the absence of *SORL1*. We also observed decreased colocalization of APP with Rab7, a marker of maturing endosomes (**Figure 2 E-F**), suggesting that in the context of *SORL1* deficiency, vesicles with APP cargo are either not maturing into late endosomes/lysosomes, or that the trafficking of APP itself to these compartments is impaired. Although the enlarged endosome morphology we observe is amyloid independent, downstream amyloidogenic cleavage of APP occurs in acidic early endosomes[31], where we document increased localization of APP. Previous studies have shown that molecules that enhance retrograde trafficking away from the early endosome towards the Golgi or back to the plasma membrane reduce amyloidogenic cleavage and decrease colocalization of APP with early endosomal markers[32, 33]. The impact of *SORL1* expression on APP processing has been previously described by us and others in studies showing knock-down of *SORL1* increases Aβ peptides in non-neuronal cells[4] and in hiPSC-derived neurons[20]. We confirmed that *SORL1* KO neurons have increased Aβ peptides released into the culture media, and that both Aβ 1-40 and Aβ 1-42 species were equally increased, without inducing a significant change in the Aβ 42:40 ratio (**Figure 2G-H**). We did not observe a change in APP holoprotein expression in *SORL1* KO neurons (**Supplemental Figure 2).** While BACE1 inhibition significantly reduced Aβ levels in all cell lines (**Figure 2G**), there were still detectable Aβ peptides in the neuronal media, showing that there was not a complete inhibition of BACE1. In concordance with an increase of APP in early endosomes in the absence of *SORL1* (**Figure 2A**), there were still significantly higher levels of Aβ peptides secreted into the culture media in the *SORL1* KO neurons, even in the presence of BACE1 inhibition (**Figure 2G**).

## Discussion

Endosome enlargement is an early cytopathology in AD[2] and abnormalities in the endo/lysosomal network are prevalent throughout human neurodegenerative disorders[34]. Multiple genetic studies also identify risk loci in or near genes involved in endosomal trafficking[35]. This points to protein trafficking as an important, and potentially modifiable, pathway for Alzheimer’s disease and related disorders. Importantly, as defects in the endosomal network are an early event in AD pathogenesis, this is an important target whose modulation will impact downstream pathologies such as Aβ and tau prior to toxic accumulations of these proteins. Indeed, our previous work has shown that molecules that enhance endosomal trafficking pathways affect Aβ and Tau independently, supporting the premise that trafficking dysfunction is an early driver of AD [33]. In this study, we used hiPSC and CRISPR/Cas9 technology to ask whether loss of an established AD risk gene, *SORL1*, induces early endosome pathology in neurons and other cell types affected in AD. Using an unbiased, quantitative confocal microscopy approach, we documented significantly enlarged endosomes in hiPSC-derived neurons lacking *SORL1*, demonstrating that loss of this sorting receptor is sufficient to induce this pathology in neurons (**Figure 1**).

Our studies confirm in human neurons that *SORL1* functions is a retromer-receptor that traffics APP out of endosomes, and that by increasing the resident time of APP in endosomal membranes *SORL1* deficiency accelerates amyloidogenic APP cleavage. Previous studies have established that cleaved fragments of APP can themselves cause endosomal enlargement[26], and so we utilized our hiPSC system to determine whether the enlargement we observed is mediated via APP cleaved products. By using BACE1 inhibitors to inhibit production of APP cleavage peptides, we show that *SORL1* deficiency still caused endosomal enlargement, showing that *SORL1* causes AD’s hallmark cytopathology in an amyloid-independent manner.

The genetics of autosomal-dominant, early-onset, AD (pathogenic variants in *APP* or *PSEN*s) belong to the same biological pathway—APP processing. In contrast, genetic studies of sporadic AD have identified approximately two dozen associated candidate genes, a group of which cohere around vesicular trafficking and endocytosis pathways. Among these, *SORL1* is common and when truncation mutations occur that cause complete loss-of-function of the SorLA protein, they are shown to be causal for late-onset AD[7, 25] making *SORL1* depletion a reliable approach for studying this pathway. Taken together with other recent findings[26], our study establishes that the two causative pathways in AD—APP cleavage and endosomal trafficking— independently cause AD’s hallmark cytopathology.

Endosome dysfunction is seen prior to Aβ accumulation in AD brains[2] and several studies have implicated the APP βCTF, rather than Aβ itself, as a possible primary toxic molecule[26, 36]. Recently, in a large cohort of FAD hiPSC-derived neurons, endosome enlargement due to FAD mutations was rescued by BACE1 inhibition[26]. In our study, BACE1 inhibition reduced Aβ peptides but did not ameliorate, and in fact exacerbated, endosomal size increases induced by loss of *SORL1* in hiPSC-derived neurons (**Figure 1**). This suggests endosomal homeostasis may require a delicate balance of cargo, and that endosome pathology is a global event in AD pathogenesis impacted by many different factors, which likely differ between early and late onset forms of the disease.

Enlarged early endosomes were first reported in neurons in post-mortem brain[2] and models using hiPSCs with FAD mutations or from SAD patients have also shown endosome enlargement in neurons[27, 36, 37]. In addition to neurons, microglia (the resident immune cells of the CNS) are also highly reliant on the endosomal network for the trafficking of internalized substrates[38]. We differentiated microglial-like cells from our *SORL1* deficient hiPSCs and performed the same analysis as with hiPSC-derived neurons. Interestingly, we did not observe significant endosome enlargement in *SORL1* deficient microglia (**Figure 1**). This data demonstrates that loss of *SORL1* impacts microglial cells differently than neuronal cells, emphasizing important differences in cell-type specific responses to insults involving the endosomal network. For example, recent work reported that loss of the endosomal adaptor protein TOM1 in microglia led to reduced microglial branching and impaired phagocytosis, while in neurons it resulted in upregulation of inflammatory signaling genes[39]. Due to the vastly different roles microglia and neurons play in the CNS, future work on analyzing the functionality of endosomal trafficking, in addition to endosome size, in *SORL1* deficient microglial-like cells is warranted.

In neurons, one of the best-characterized cargos of *SORL1* is APP. The protein SorLA directly binds APP[40] and serves as an adaptor molecule, via VPS26, in retromer-dependent retrograde trafficking of APP[18]. Because APP localization within the endosomal network dictates amyloidogenic processing of APP, we performed colocalization analysis of APP with early endosomes, late endosomes, and the trans-Golgi network to determine how loss of *SORL1* may impact APP subcellular localization in human neurons. We observed that loss of *SORL1* alters APP trafficking in the endosomal network, leading to increased colocalization of APP in early endosomes and a decrease in its localization in retrograde pathways and the TGN. This is in line with studies in other model systems[18, 32] and confirms that *SORL1* plays an important role in regulating APP amyloidogenic cleavage. We also observe that APP localization is decreased in Rab7+ late endosomes/lysosomes, suggesting a defect in either endosomal maturation or trafficking of APP towards degradative compartments (**Figure 2**).

Taken together, our work confirms the importance of *SORL1* in preventing amyloidogenic processing of APP, and suggests that it may have a broader role in regulating endosomal network function. Our data supports the idea that endosome pathology is an upstream event from Aβ generation and suggests that further disruptions of endosomal cargo processing or homeostasis (such as BACE inhibition) can enhance traffic jams or delay maturation in the absence of *SORL1*. Finally, our work demonstrates that hiPSC-derived neuronal models are valuable tools for dissecting early pathogenic events in AD, and may help to mechanistically clarify the molecular mechanisms that underlie therapeutic failures.

## Acknowledgements

We would like to thank the members of the Young laboratory for crucial discussions in the preparation of this manuscript and Dr. Gregory Petsko for critical feedback on the manuscript. We would like to acknowledge the UW SLU Cell Analysis Facility and the Garvey Imaging Core at the UW Institute for Stem Cell and Regenerative Medicine. This work was supported by NIH grants (R01AG062148) and BrightFocus grant (A2018656S) to JEY, a Biogen Sponsored Research Agreement to JEY, a NIH Training grant (T32AG052354) to AK and a generous gift from the Ellison Foundation (to UW).

## Author Contributions

AK, SM, DWH, SAS, and JEY conceptualized and designed the experiments. AK, SM, RM and MS performed the experiments. AK, SM and JEY analyzed the data. JB and SJ differentiated microglial-like cells from hiPSCs. JEY conceived the project and wrote the manuscript with assistance and editing from AK, SM, SAS, and SJ.

## Supplemental Figure legends

**Supplemental Figure 1:**
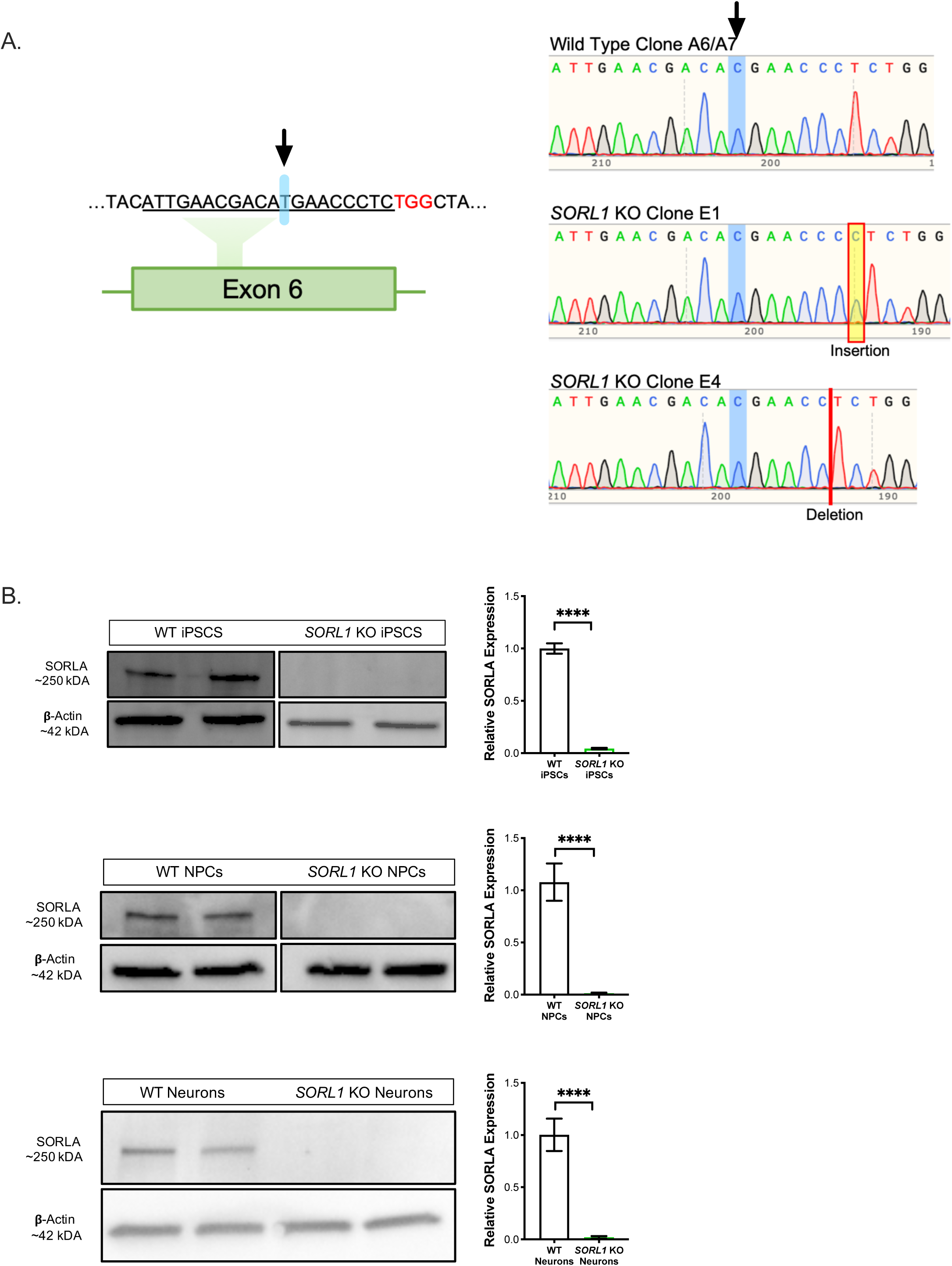
A. Schematic drawing representing CRISPR/Cas9 depletion of *SORL1* in hiPSCs. The gRNA (underlined) was targeted to exon 6 in the *SORL1* coding sequence. The light blue shading, indicated by an arrowhead, indicates a synonymous SNP present in the donor hiPSC line that differs from the reference genome. The two KO clones used in this study showed and insertion (E1) and a deletion (E4) three bases upstream of the PAM site that leads to a frameshift and premature stop codon. B. Representative Western blots and quantitation show reduction of SORLA protein levels to nearly zero in the KO hiPSCs, NPCs, and neurons. Quantification for all cell types, n=6 biological replicates. All values represent mean +/-SD. All normally distributed data were analyzed by two-tailed unpaired t-test *p<.05, **p<.01, ***p<.001, ****p<.0001.

**Supplemental Figure 2:**
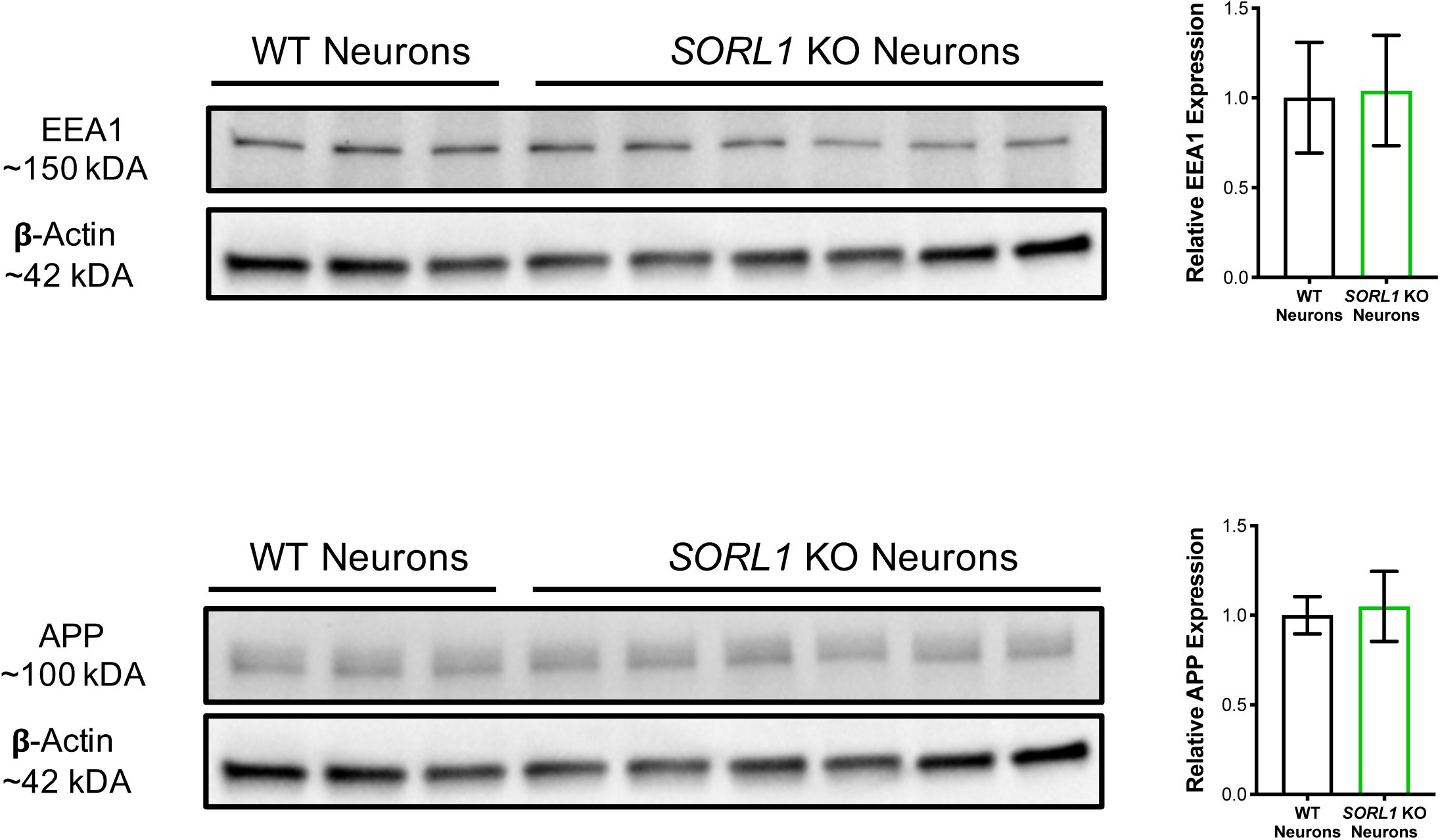
A. *SORL1* KO neurons do not have significant differences in EEA protein expression. B. *SORL1* KO neurons do not have significant differences in full-length APP expression. N= 3-6 biological replicates. All values represent mean +/-SD. All normally distributed data were analyzed by two-tailed unpaired t-test *p<.05, **p<.01, ***p<.001, ****p<.0001.

**Supplemental Figure 3:**
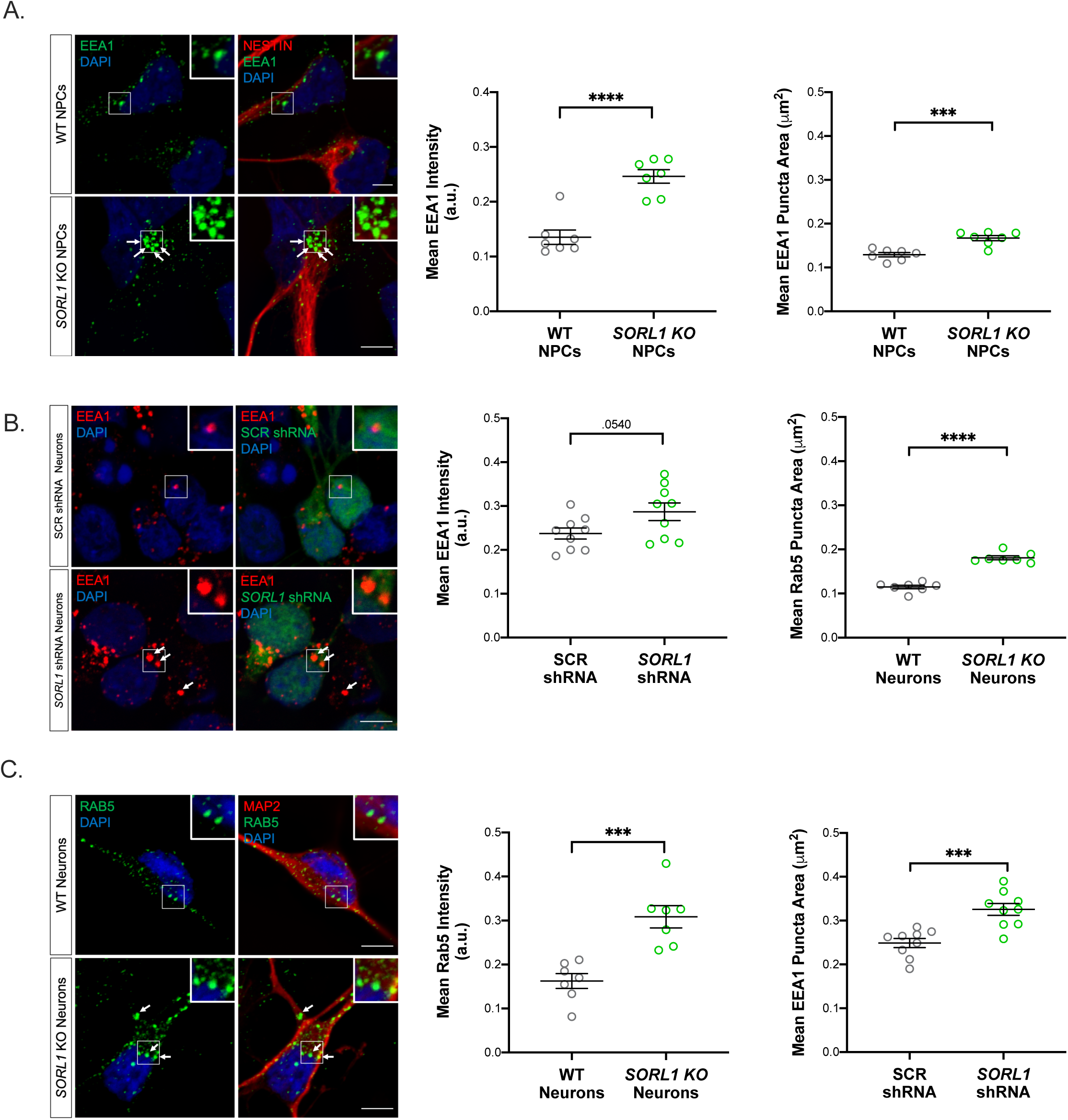
A. Representative immunofluorescence images and quantitation of enlarged EEA positive early endosomes in *SORL1* KO and isogenic WT neural progenitor cells (NPCs). N=7 images (24-26 cells) B. Representative immunofluorescence images and quantitation of enlarged EEA1 positive early endosomes in WT hiPSC-derived neurons treated with a *SORL1* shRNA. N=9 images (36-48 cells) C. Representative immunofluorescence images and quantitation of enlarged Rab5 positive early endosomes in *SORL1* KO and isogenic WT hiPSC-derived neurons. N= 9 images (36-48 cells). All values represent mean +/- SEM. *p<.05, **p<.01, ***p<.001, ****p<.0001 by two-tailed, unpaired T-test.

**Supplemental Figure 4:**
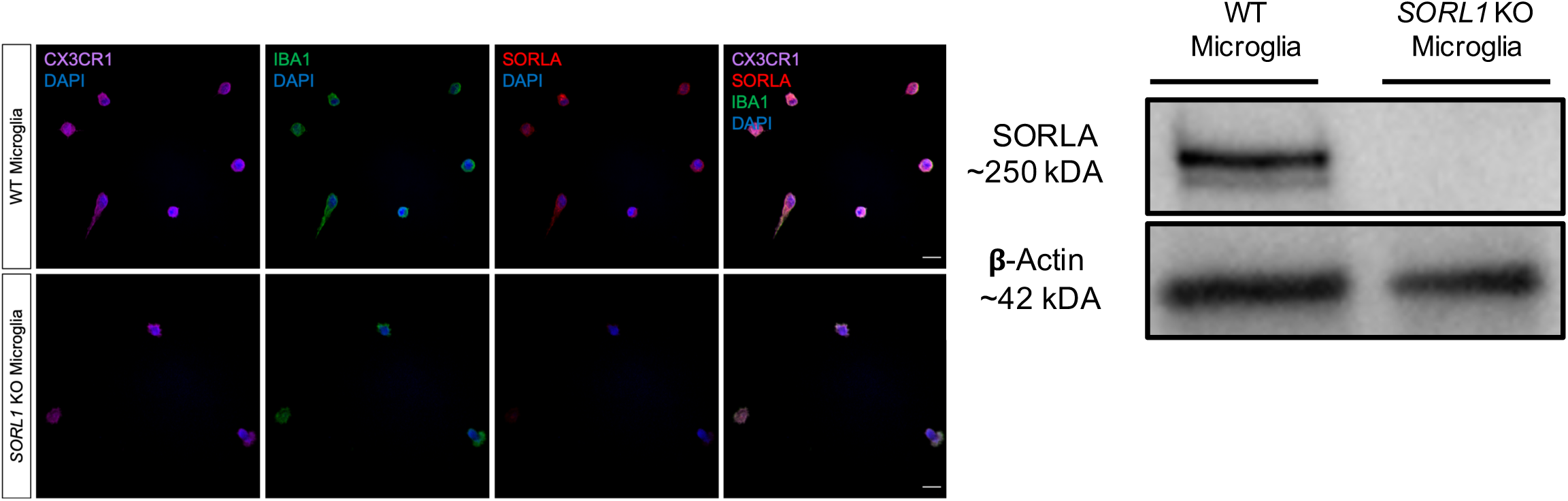
Representative immunofluorescence images and Western blot of *SORL1* KO and isogenic wild-type microglial-like cells differentiated from hiPSCs. Microglia-like cells show expression of microglial markers IBA1 and CX3CR1. *SORL1* KO microglil-like cells show no SorLA staining and no SorLA expression by Western blot. Scale bars are 20μm.

## LEAD CONTACT AND MATERIALS AVAILABILITY

Further information and requests for resources and reagents should be directed to and will be fulfilled by the Lead Contact, Jessica Young.

## EXPERIMENTAL MODEL AND SUBJECT DETAILS

### CELL LINES

#### CRISPR/Cas9 Genome Editing

All genome editing was performed in the previously published and characterized male CV background line[20]. Genome edited lines were generated using published protocols[33]. Briefly, guide RNAs (gRNAs) to SORL1 were generated using the Zhang Lab CRISPR Design website at MIT (crispr.mit.edu) and selected to minimize off-target effects. gRNAs were cloned into vector px458 that co-expresses the Cas9 nuclease and GFP, and hiPSCs were electroporated with the plasmid. Electroporated hiPSCs were FACS sorted for GFP, plated in 10cm plates at a clonal density (∼1×10^4 cells/plate), and allowed to grow for roughly 2 weeks. Colonies were picked into 96 well plates and split into two identical sets. One set was analyzed for sequence information by Sanger sequencing and one set was expanded for cell line generation.

#### CRISPR/Cas9 gRNA, ssODN, and Primer Sequences

gRNA: ATTGAACGACATGAACCCTC

ssOD: GGGAATTGATCCCTATGACAAACCAAATACCATCTACATTGAACGACATGAACCCT CTGGCTACTCCACGTCTTCCGAAGTACAGATTTCTTCCAGTCCCGGGAAAACCAGG AAG

Forward primer: ctctatcctgagtcaaggagtaac

Reverse primer: ccttccaattcctgtgtatgc

PCR amplifies 458 bp sequence

#### Cell Line Karyotyping

Karyotyping analysis was performed on hiPSC lines by Diagnostic Cytogenetics, Inc. (Seattle, WA).

## METHOD DETAILS

### Western Blotting

Cell lysates were run on BioRad 4-10% Mini-PROTEAN TGX Precast Protein Gels (BioRad, Hercules, CA) and transferred to PVDF membranes. Membranes were probed with antibodies to Sortilin-related receptor 1 (SORLA) at 1:1000 (BD 611860 and abcam ab190684), β-actin (ACTB) at 1:2000 (EMD Millipore Corp MAB1501), early endosome antigen 1 (EEA1) at 1:5000 (BD 610456), and amyloid precursor protein (APP) at 1:500 (Invitrogen 14-9749-80). Imaging was performed with a BioRad ChemiDoc system and quantification was performed using ImageJ software.

### hiPSC Neuronal Differentiation

hiPSCs were differentiated to neurons using previously described dual-SMAD inhibition techniques[41]. Briefly, hiPSCs were plated on Matrigel coated 6-well plates at a density of 3.5 million cells per well and fed with Basal Neural Maintenance Media (1:1 DMEM/F12 + glutamine media/neurobasal media, 0.5% N2 supplement, 1% B27 supplement, 0.5% GlutaMax, 0.5% insulin-transferrin-selenium, 0.5% NEAA, 0.2% β-mercaptoethanol; Gibco, Waltham, MA) + 10μM SB-431542 + 0.5μM LDN-193189 (Biogems, Westlake Village, CA). Cells were fed daily for seven days. On day eight, cells were incubated with Versene, gently dissociated using cell scrapers, and split 1:3. On day nine media was switched to Basal Neural Maintenance Media + 20 ng/mL FGF (R&D Systems, Minneapolis, MN) and fed daily. On day sixteen, cells were passaged 1:3. Cells were fed until approximately day twenty-three. At this time, cells were FACS sorted to obtain the CD184/CD24 positive, CD44/CD271 negative neural precursor cell (NPC) population[42]. Following sorting, NPCs were expanded for neural differentiation. For cortical neuronal differentiation, NPCs were plated out in 10cm plates at a density of 6 million cells/plate. After 24 hours, cells were switched to Neural Differentiation media (DMEM-F12 + glutamine, 0.5% N2 supplement, 1% B27 supplement, 0.5% GlutaMax) + 0.2μg/mL brain-derived neurotrophic factor (PeproTech, Rocky Hill, NJ) + 0.2μg/mL glial-cell-derived neurotrophic factor (PeproTech) + 0.5M dbcAMP (Sigma Aldrich, St Louis, MO). Media was refreshed twice a week for three weeks. After three weeks, neurons were selected for CD184/CD44 negative population by MACS sorting and plated for experiments.

### Purification of Neurons

Following three weeks of differentiation, cells were dissociated with accutase and resuspended in IMAG solution (PBS + 0.5% bovine serum albumin [Sigma Aldrich, St Louis, MO] + 2mM ethylenediaminetetraacetic acid [ThermoFisher Scientific, Waltham, MA]). Following a modification of Yuan et al., 2011[42], cells were incubated with PE-conjugated mouse anti-Human CD44 and mouse anti-Human CD184 antibodies (BD Biosciences, San Jose, CA) at a concentration of 5μL/10 million cells. Following antibody incubation, cells were washed with IMAG solution and incubated with anti-PE magnetic beads (BD Biosciences, San Jose, CA) at a concentration of 25μL/10 million cells. Bead-antibody complexes were pulled down using a rare earth magnet, supernatants were selected, washed, and plated at an appropriate density.

### Differentiation of iPSCs into microglia

SORL1 KO and WT iPSCs were differentiated into microglia as previously described[29]. Briefly, iPSCs were plated in mTESR™ 1 medium (#05850; STEMCELL Technologies) supplemented with ROCK Inhibitor (Y-27632; # A3008; Apex Bio) on Matrigel (Growth factor reduced basement membrane matrix; # 356231; Corning) coated 6 well plates (#657160; CELLSTAR) at a dilution of 1:30. To begin hematopoietic progenitor differentiation, these cells were passaged to get a density of ∼100 colonies (∼150 cells per colony of iPSCs) per well of a 6 well plate. On day 0, mTESR™ 1 medium was replaced with STEMdiff™ Hematopoietic Supplement A medium from the STEMdiff™ Hematopoietic kit (# 05310; STEMCELL technologies). On day 3, when colonies became flattened, medium was replaced with STEMdiff™ Hematopoietic Supplement B medium from the STEMdiff™ Hematopoietic kit (# 05310; STEMCELL technologies). Cells remained in this medium for 7 additional days. By day 10, non-adherent hematopoietic progenitor cells (HPCs) coming off from the flattened colonies were harvested by removing medium. Any remaining HPCs/floating cells were collected by gentle PBS washes. At this point, HPCs were either frozen using Bambanker cell freezing medium (#BBH01; Bulldog-Bio) or plated at a density of 0.2M cells per well of a Matrigel coated (1:60 dilution) 6 well plate in microglia differentiation medium for 25 days. Microglia differentiation medium comprised of DMEM-F12 (#11039047; Thermo Fisher Scientific), Insulin-transferrin-selenite (#41400045; Thermo Fisher Scientific), B27 (# 17504-044; Thermo Fisher Scientific), N2 (# 17502-048; Thermo Fisher Scientific), glutamax (# 35050061; Thermo Fisher Scientific), non-essential amino acids (# 11140050; Thermo Fisher Scientific), monothioglycerol (# M1753; Sigma), Insulin (# I2643; Sigma) freshly supplemented with TGF-β (#130-108-969, Miltenyl), IL-34 (# 200-34; Peprotech) and M-CSF (#PHC9501; Thermo Fisher Scientific). On day 25, this medium was supplemented with CD200 (#C311; Novoprotein) and CX3CL1 (#300-31; Peprotech) for maturation of microglia. Cells remained in this medium for 3 days. On day 28, microglia differentiation was complete, and these cells were plated in laminin (#L2020; Sigma) coated coverslips (12mm diameter, #1760-012; cglifesciences) in a 24 well plate for immunocytochemistry with appropriate antibodies.

### Amyloid Beta measurements

Aβ peptides and phosphorylated tau were measured as previously described (Young 2015). Briefly, purified neurons were seeded at a density of 200,000 cells/well of a 96-well plate and maintained in culture for 5 days. Medium and lysates were harvested from triplicate wells. To measure secreted Aβ peptides, media was run on an Aβ Triplex ELISA plate (Meso Scale Discovery).

### Immunocytochemistry

Purified neurons were seeded at a density of 500,000 cells per well of a 24-well plate on glass coverslips coated with Matrigel. After 5 days in culture, cells were fixed in 4% paraformaldehyde (PFA, Alfa Aesar, Reston, VA) for 15 minutes. Cells were incubated in blocking buffer containing 2.5% bovine serum albumin and 0.1% Triton X-100 (Sigma Aldrich, St Louis, MO) for 30 minutes at room temperature then incubated in a primary antibody dilution in blocking buffer for 2 hours at room temperature. Cells were washed 3x with PBS + 0.1% Triton X-100 and incubated with a secondary antibody dilution in blocking buffer for 1 hour at room temperature. Cells were washed 3x in PBS and mounted on glass slides with ProLong Gold Antifade mountant (ThermoFisher, Waltham, MA). The following primary antibodies were used: Ras-related protein Rab-5A (RAB5A) at 1:500 (Synaptic Systems 108 011); early endosome antigen 1 (EEA1) at 1:500 (BD 610456); amyloid precursor protein (APP) at 1:250 (Abcam ab32136); microtubule-associated protein 2 (Map2) at 1:1000 (Abcam ab92434); Nestin (NES) at 1:1000 (Santa Cruz Biotechnology sc23927); Trans-Golgi network integral membrane protein (TGN38) at 1:250 (Santa Cruz sc-166594); Ras-related protein Rab-7a (Rab7) at 1:1000 (Abcam ab50533); DAPI at 1 μg/mL final (Alfa Aesar).

### Confocal Microscopy and Image Processing

All microscopy and image processing were performed under blinded conditions. Confocal z-stacks were obtained using a Nikon A1R confocal microscope with x63 and x100 plan apochromat oil immersion objectives. Image processing was performed using ImageJ software. For endosome analysis, maximum intensity projections of confocal stacks were generated, and background was subtracted using the rolling ball algorithm. Endosome channels were enhanced using contrast limited adaptive histogram equalization algorithms (CLAHE) and masked using cell body stains. Size and intensity measurements were performed using CellProfiler software. Individual puncta were identified using automated segmentation algorithms. Mean intensity of each puncta was measured and has been presented as a mean value over all puncta per field. Similarly, pixel area of each puncta was measured and has been presented as a mean area over all puncta per field. For colocalization analysis, a minimum of 10 fields of confocal z-stacks were captured using the analyzed using the x100 plan apochromat oil immersion objective. Median filtering was used to remove noise from images and manual thresholding was applied to all images. The colocalization of APP with endocytic markers was quantified using JACOP plugin in Image J and presented as Mander’s correlation coefficient.

### BACE Inhibition

Purified neurons derived from SORL1 KO and WT iPSCs were plated on a matrigel coated 96 well plate at a density of 2×10^5^ cells per well. After 5 days, cells were treated with either 25nM β-secretase inhibitor (BACEi; LY2886721; # HY-13240; MedChemExpress) or DMSO (as a vehicle control) for 72h. All experiments were performed after 72h of drug treatment. At this point, medium from DMSO or BACEi treated neurons was harvested for quantification of Aβ-40 and Aβ-42 peptides secreted by neurons by ELISA. Additionally, cell lysates were harvested for determining protein levels of β-C terminal fragment (β-CTF) of Amyloid Precursor Protein (APP) by western blot.

## QUANTIFICATION AND STATISTICAL ANALYSIS

We used two independent clones of homozygous knockout cell lines and two independent clones of isogenic wild-type cell lines (cells that underwent the CRISPR/Cas9 transfection and sub-cloning process, but in which editing events did not occur). Experimental data was tested for normal distributions. Normally distributed data was analyzed using parametric ANOVA tests and non-normally distributed data was analyzed by non-parametric Kruskal-Wallis tests using GraphPad Prism software. For imaging analysis, 10-20 fields were analyzed for a total of 10-58 cells. Statistical details of individual experiments can be found in figure legends.

## DATA AND CODE AVAILABILITY

No datasets or code were generated during this study. The images and CellProfiler pipelines used in this study are available from the corresponding author on request.

## References

1. Alzheimer’s Association Report: 2019 Alzheimer’s disease facts and figures. Alzheimer’s & Dementia, 2019. 15: p. 321–387.

2. Cataldo, A.M., et al., Endocytic pathway abnormalities precede amyloid beta deposition in sporadic Alzheimer’s disease and Down syndrome: differential effects of APOE genotype and presenilin mutations. Am J Pathol, 2000. 157(1): p. 277–86.

3. Offe, K., et al., The lipoprotein receptor LR11 regulates amyloid beta production and amyloid precursor protein traffic in endosomal compartments. J Neurosci, 2006. 26(5): p. 1596–603.

4. Rogaeva, E., et al., The neuronal sortilin-related receptor SORL1 is genetically associated with Alzheimer disease. Nat Genet, 2007. 39(2): p. 168–77.

5. Karch, C.M. and A.M. Goate, Alzheimer’s disease risk genes and mechanisms of disease pathogenesis. Biol Psychiatry, 2015. 77(1): p. 43–51.

6. Bettens, K., et al., SORL1 is genetically associated with increased risk for late-onset Alzheimer disease in the Belgian population. Human Mutation, 2008. 29(5): p. 769–770.

7. Holstege, H., et al., Characterization of pathogenic SORL1 genetic variants for association with Alzheimer’s disease: a clinical interpretation strategy. Eur J Hum Genet, 2017. 25(8): p. 973–981.

8. Pottier, C., et al., High frequency of potentially pathogenic SORL1 mutations in autosomal dominant early-onset Alzheimer disease. Mol Psychiatry, 2012. 17(9): p. 875–9.

9. Reitz, C., et al., Meta-analysis of the association between variants in SORL1 and Alzheimer disease. Arch Neurol, 2011. 68(1): p. 99–106.

10. Andersen, O.M., et al., Neuronal sorting protein-related receptor sorLA/LR11 regulates processing of the amyloid precursor protein. Proc Natl Acad Sci U S A, 2005. 102(38): p. 13461–6.

11. Hermans-Borgmeyer, I., et al., Unique expression pattern of a novel mosaic receptor in the developing cerebral cortex. Mech Dev, 1998. 70(1-2): p. 65–76.

12. Zhang, Y., et al., An RNA-sequencing transcriptome and splicing database of glia, neurons, and vascular cells of the cerebral cortex. J Neurosci, 2014. 34(36): p. 11929–47.

13. Dumanis, S.B., et al., Distinct Functions for Anterograde and Retrograde Sorting of SORLA in Amyloidogenic Processes in the Brain. J Neurosci, 2015. 35(37): p. 12703–13.

14. Glerup, S., et al., SorLA controls neurotrophic activity by sorting of GDNF and its receptors GFRalpha1 and RET. Cell Rep, 2013. 3(1): p. 186–99.

15. Herskowitz, J.H., et al., GGA1-mediated endocytic traffic of LR11/SorLA alters APP intracellular distribution and amyloid-beta production. Mol Biol Cell, 2012. 23(14): p. 2645–57.

16. Nielsen, M.S., et al., Sorting by the cytoplasmic domain of the amyloid precursor protein binding receptor SorLA. Mol Cell Biol, 2007. 27(19): p. 6842–51.

17. Klinger, S.C., et al., SorLA regulates the activity of lipoprotein lipase by intracellular trafficking. J Cell Sci, 2011. 124(Pt 7): p. 1095–105.

18. Fjorback, A.W., et al., Retromer binds the FANSHY sorting motif in SorLA to regulate amyloid precursor protein sorting and processing. J Neurosci, 2012. 32(4): p. 1467–80.

19. Mehmedbasic, A., et al., SorLA complement-type repeat domains protect the amyloid precursor protein against processing. J Biol Chem, 2015. 290(6): p. 3359–76.

20. Young, J.E., et al., Elucidating Molecular Phenotypes Caused by the SORL1 Alzheimer’s Disease Genetic Risk Factor Using Human Induced Pluripotent Stem Cells. Cell Stem Cell, 2015. 16(4): p. 373–85.

21. Dodson, S.E., et al., LR11/SorLA expression is reduced in sporadic Alzheimer disease but not in familial Alzheimer disease. J Neuropathol Exp Neurol, 2006. 65(9): p. 866–72.

22. Ma, Q.L., et al., Reduction of SorLA/LR11, a sorting protein limiting beta-amyloid production, in Alzheimer disease cerebrospinal fluid. Arch Neurol, 2009. 66(4): p. 448–57.

23. Sager, K.L., et al., Neuronal LR11/sorLA expression is reduced in mild cognitive impairment. Ann Neurol, 2007. 62(6): p. 640–7.

24. Caglayan, S., et al., Lysosomal sorting of amyloid-beta by the SORLA receptor is impaired by a familial Alzheimer’s disease mutation. Sci Transl Med, 2014. 6(223): p. 223ra20.

25. Raghavan, N.S., et al., Whole-exome sequencing in 20,197 persons for rare variants in Alzheimer’s disease. Ann Clin Transl Neurol, 2018. 5(7): p. 832–842.

26. Kwart, D., et al., A Large Panel of Isogenic APP and PSEN1 Mutant Human iPSC Neurons Reveals Shared Endosomal Abnormalities Mediated by APP beta-CTFs, Not Abeta. Neuron, 2019. 104(5): p. 1022.

27. Lin, Y.T., et al., APOE4 Causes Widespread Molecular and Cellular Alterations Associated with Alzheimer’s Disease Phenotypes in Human iPSC-Derived Brain Cell Types. Neuron, 2018.

28. Wang, C., et al., Gain of toxic apolipoprotein E4 effects in human iPSC-derived neurons is ameliorated by a small-molecule structure corrector. Nat Med, 2018. 24(5): p. 647–657.

29. McQuade, A., et al., Development and validation of a simplified method to generate human microglia from pluripotent stem cells. Mol Neurodegener, 2018. 13(1): p. 67.

30. Kaur, G. and A. Lakkaraju, Early Endosome Morphology in Health and Disease. Adv Exp Med Biol, 2018. 1074: p. 335–343.

31. Small, S.A. and S. Gandy, Sorting through the cell biology of Alzheimer’s disease: intracellular pathways to pathogenesis. Neuron, 2006. 52(1): p. 15–31.

32. Mecozzi, V.J., et al., Pharmacological chaperones stabilize retromer to limit APP processing. Nat Chem Biol, 2014. 10(6): p. 443–9.

33. Young, J.E., et al., Stabilizing the Retromer Complex in a Human Stem Cell Model of Alzheimer’s Disease Reduces TAU Phosphorylation Independently of Amyloid Precursor Protein. Stem Cell Reports, 2018. 10(3): p. 1046–1058.

34. Vagnozzi, A.N. and D. Pratico, Endosomal sorting and trafficking, the retromer complex and neurodegeneration. Mol Psychiatry, 2019. 24(6): p. 857–868.

35. Van Acker, Z.P., M. Bretou, and W. Annaert, Endo-lysosomal dysregulations and late-onset Alzheimer’s disease: impact of genetic risk factors. Mol Neurodegener, 2019. 14(1): p. 20.

36. Israel, M.A., et al., Probing sporadic and familial Alzheimer’s disease using induced pluripotent stem cells. Nature, 2012. 482(7384): p. 216–20.

37. Raja, W.K., et al., Self-Organizing 3D Human Neural Tissue Derived from Induced Pluripotent Stem Cells Recapitulate Alzheimer’s Disease Phenotypes. PLoS One, 2016. 11(9): p. e0161969.

38. Sole-Domenech, S., et al., The endocytic pathway in microglia during health, aging and Alzheimer’s disease. Ageing Res Rev, 2016. 32: p. 89–103.

39. Martini, A.C., et al., Amyloid-beta impairs TOM1-mediated IL-1R1 signaling. Proc Natl Acad Sci U S A, 2019. 116(42): p. 21198–21206.

40. Andersen, O.M., et al., Molecular dissection of the interaction between amyloid precursor protein and its neuronal trafficking receptor SorLA/LR11. Biochemistry, 2006. 45(8): p. 2618–28.

41. Rose, S.E., et al., Leptomeninges-Derived Induced Pluripotent Stem Cells and Directly Converted Neurons From Autopsy Cases With Varying Neuropathologic Backgrounds. J Neuropathol Exp Neurol, 2018.

42. Yuan, S.H., et al., Cell-surface marker signatures for the isolation of neural stem cells, glia and neurons derived from human pluripotent stem cells. PLoS One, 2011. 6(3): p. e17540.

